# I judge you by your profit: Judgments of effort exerted by self and others are influenced by received rewards

**DOI:** 10.1101/253138

**Authors:** Max Rollwage, Franziska Pannach, Caedyn Stinson, Ulf Toelch, Igor Kagan, Arezoo Pooresmaeili

**Author notes:** Correspondences should be addressed to:, Max Rollwage. IK and AP are joint last authors.

## Abstract

Effort constitutes a major part of cost-benefit calculations underlying decision making. Therefore, estimating the effort someone has spent on a task is a core dimension for evaluating own and others’ actions. It has been previously shown that self-judgments of effort are influenced by the magnitude of obtained rewards. It is unclear, however, whether the influence of reward on effort estimations is limited to self-judgments or whether reward incorporation represents a general computational principle when judging effort. Here we show that people also integrate reward magnitude when judging the effort exerted by others. Participants (N=48) performed an effortful sensorimotor task interleaved with a partner, while rating either their own or the other person’s effort. After each trial but before the effort rating, both participants were informed about the obtained reward. We found that higher rewards led to higher estimations of exerted effort, in self-as well as other-judgments, and this effect was more pronounced for other-judgments. In both types of judgment, computational modelling revealed that reward information and the perceived level of exertion were combined in a Bayes optimal manner to form effort estimates. Remarkably, the extent to which rewards influenced effort judgments was positively correlated with conservative world-views, indicating that the basic computations underlying this behavioural phenomenon might be related to more general beliefs about the association between effort and reward in the society. The integration of reward information into retrospective effort judgments underscores the convergence of multiple information sources that supports adaptive learning and decision making in social contexts.

## Introduction

Every action comes at a cost and we constantly evaluate in a retrospective manner whether an effortful behaviour was “worth it”. Effort judgments are particularly important in a social context as effort is one of the core dimensions for evaluating behaviour of others. For instance, people that contributed more effort to a cooperative task are perceived as more likeable (1), while freeriders, who do not contribute to the joint endeavour, are punished (2, 3). Outside the lab, effort estimations influence group dynamics and broader social attitudes. For instance the support for welfare benefits is strongly linked to the perception that the people who receive those benefits have made an effort to find a job (4). Although retrospective effort estimations have an important impact on different aspects of our lives, decision neuroscience has so far mainly focused on prospective effects of expected effort on action plans. The common finding of these studies is that effort is processed as a value discounting factor that is traded off against potential rewards when making choices (5-11). However, decision making often goes beyond an immediate comparison between costs and benefits and involves evaluating a stream of actions and outcomes that occur over time (12). As such, effort judgments at any given moment inherently involve a retrospective evaluation of the exerted effort in the past and the resultant individual or social outcomes. Surprisingly, the underlying mechanisms of retrospective evaluations of effort and their relationship to the decisions pertaining to future costs of actions have been underexplored in cognitive neurosciences. In comparison, social psychology has extensively investigated retrospective evaluations of effort and interactions between attribution of rewards and effort depending on the social context (e.g. attribution theory (13) and effort justification (14)). For instance, it has been shown that people differ in the attribution of success when evaluating their own behaviour (where success is attributed to more effort) or others’ behaviour (where success is attributed to luck (13)). However, to understand the underlying mechanisms of such interactions during decision making, a computational approach where the contributions of rewards and effort to the outcome evaluations are analytically modelled is required (15, 16).

A recent study began to fill this gap by using a computational approach to investigate different contributing factors that influence retrospective effort evaluations (17). Intuitively, people might have the impression they hold accurate representations about the effort they, and others, have spent. In contrast to this intuition, in this study self-judgments of effort were influenced by the magnitude of obtained rewards: the same level of effort was rated differently depending on the reward, reflecting a bias in effort estimations. However, it is not known whether the influence of reward on effort estimations is limited to self-judgments (i.e. representing a form of self-serving bias (18)) or whether reward information is invariably incorporated when judging exerted effort also in other contexts.

Interestingly, in the study of Pooresmaeili, Wannig and Dolan (17) the integration of rewards into effort judgments depended on the contingency between reward and task difficulty, with a higher influence of reward when it had a strong association with the task difficulty, and hence provided reliable probabilistic information about the exerted effort. Thus, the integration of reward into effort ratings resembled Bayesian cue integration (19, 20), according to which different sources of information are combined, weighted by their reliability, to infer the “true” state of the world. In such a paradigm, the consideration of rewards provides an additional source of information for estimating the exerted effort, rather than an irrational bias. If this interpretation holds true, influences of rewards on effort estimations should not be limited to self-judgments but should also be present for other-judgments, potentially showing an even stronger effect size as people ought to have less reliable internal representations for the effort exerted by others compared to their own effort.

The weighting of different information sources by their reliability might be further influenced by prior beliefs about the relevance of each information source. For instance, there is a wide range of individual differences regarding assumptions about the relationship between effort and success/reward (21, 22). The belief that success/reward is mainly driven by high effort-investment might lead to justifying economic hierarchies and contribute to stabilization of existing inequalities (17, 23, 24). Such beliefs and their ensuing attitudes show a considerable overlap with political conservatism as it has been proposed that preference for social stability and acceptance of social inequalities are the two core aspects of conservative world-views (21, 22). We therefore hypothesized that the extent to which reward influences effort estimations in a simple task might be related to broader conservative attitudes.

## Results

We investigated whether people integrate reward magnitude when judging the effort another person has exerted. We present the data of 48 subjects (24 pairs) performing an effortful task, interleaved with a partner, and rating either their own or the partner’s exerted effort. The task, adapted from Pooresmaeili, Wannig and Dolan (17), consisted of pressing two keyboard buttons as fast as possible to push a ball up a virtual ramp (**Figure 1A**). The ball rolled back with a constant displacement per time frame, simulating a gravity force. The difficulty of a given trial was determined by the gravity force (6 difficulty levels that were pseudo-randomly presented). The exact gravity force of each trial was determined by the timing of the presentation software and could slightly vary around the mean gravity for that difficulty level (**Supplementary Information**). Subjects were informed about the fact that the task contained different difficulty levels represented by varying gravity force. On trials in which the ball was successfully pushed up the ramp, the participant who conducted the trial was rewarded. The rewards were contingent on the task difficulty (with values drawn from six Gaussian distributions with means corresponding to the relative difficulty, from 1.5 to 6.5 cents, and SD 1.2), but crucially, within each difficulty level the reward was randomly selected from the corresponding distribution. In most of the trials (70%), the reward was presented before subjects rated their effort (as illustrated in **Figure 1A**) but in some of the trials (30%) the reward was presented after participants made effort judgments. Trials in which reward was shown after the effort ratings served as a reference, since in those trials no influence of reward on effort judgments should be present. In half of the trials, participants performed the task themselves and rated their own effort (self-judgment). In the other half, participants rated the other person’s effort (other-judgment) after watching them perform the task. On both self- and other-judgment trials participants observed on their screen how the ball moved up the virtual ramp and additionally heard a beep tone that corresponded to every key press through headphones (**Figure 1B**). Thus, participants received visual and auditory information about the task difficulty that should reflect the exerted physical effort of the other player.

**Figure 1.**
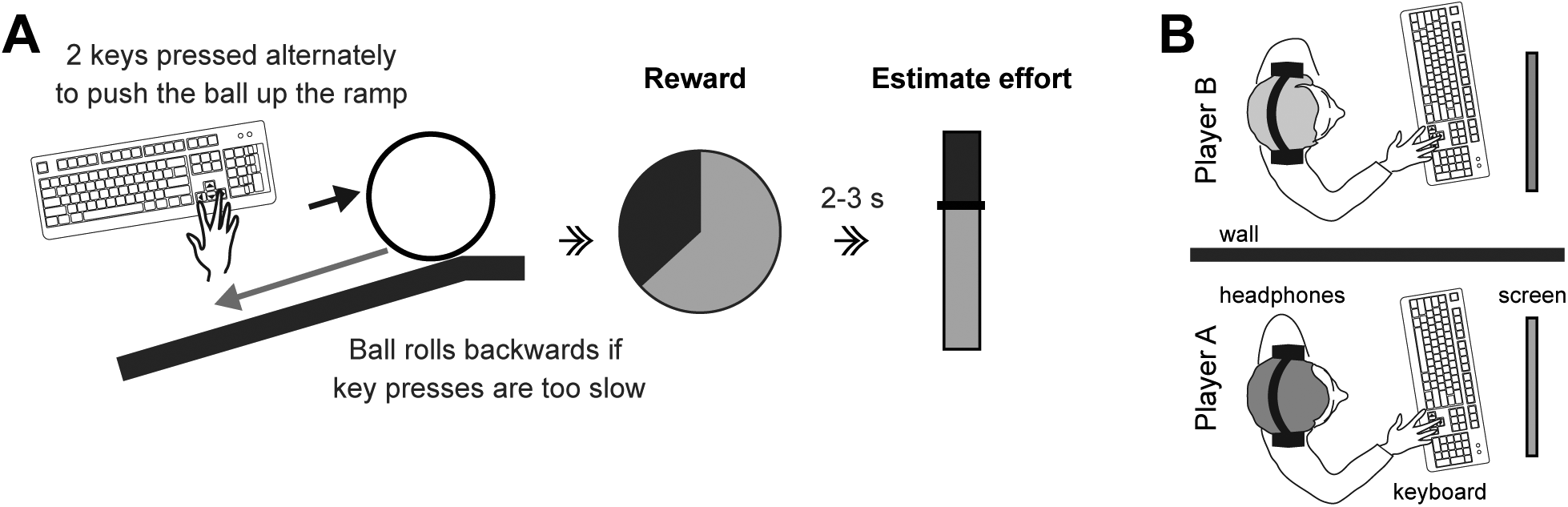
Experimental setup and task. **(A)** Participants were asked to move a ball up a ramp by performing fast, alternating key presses. A gravity force was simulated, displacing the ball backward by a constant amount on each display frame. We used six levels of task difficulty, corresponding to the amount of ball displacement per frame. After the ball was successfully pushed all the way to the top of the ramp, participants received a monetary reward, where reward amount was contingent upon task difficulty. Subjects rated their effort by shifting the position of a vertical sliding bar. In half of the trials participants performed the task themselves and rated their own effort. In the other half, participants rated the other person’s effort after watching them performing the task. **(B)** Subjects were seated next to each other in the same room, separated by a partition wall. When the partner was pressing keys to push the ball subjects saw on their screen how the ball was moved up the ramp and heard via the headphones a beep tone for every key press the other person was conducting.

We first investigated the influence of reward magnitude on effort estimations by regression analysis of trial-to-trial variation in reward magnitude and effort estimation. The analysis was conducted separately for the self- and the other-judgments of each participant and only trials in which reward information was presented before the effort rating were considered. The regression slopes were calculated between the trial-to-trial variation of reward (rewards were z-scored within each difficulty level) and estimated effort (effort ratings were z-scored within each difficulty level).

Higher rewards were associated with higher effort ratings (**Figure 2A,B**), everything else being constant. Across subjects, there was a significant influence of reward variation on effort estimation for self-judgments (one-sample signed test, p =0.002, **Figure 2C**) as well as for other-judgments (one-sample signed test, p =0.00002, **Figure 2C**). Interestingly, the effect of rewards on effort estimations was significantly stronger for other-than self-judgments (mean slopes of 0.14 and 0.1 respectively, Wilcoxon signed rank test, p =0.029, **Figure 2C**). Nevertheless, the amount of reward integration for self- and other-judgments was highly correlated across subjects (r=0.77, p<10^-11^; **Suppl. Figure S1**), indicating that integrating reward information into effort judgment is a general tendency of individuals, independent of the type of required judgment. We obtained similar results when we controlled for slight differences between actual gravity force of trials that belonged to the same difficulty level (**Supplementary Information**).

**Figure 2.**
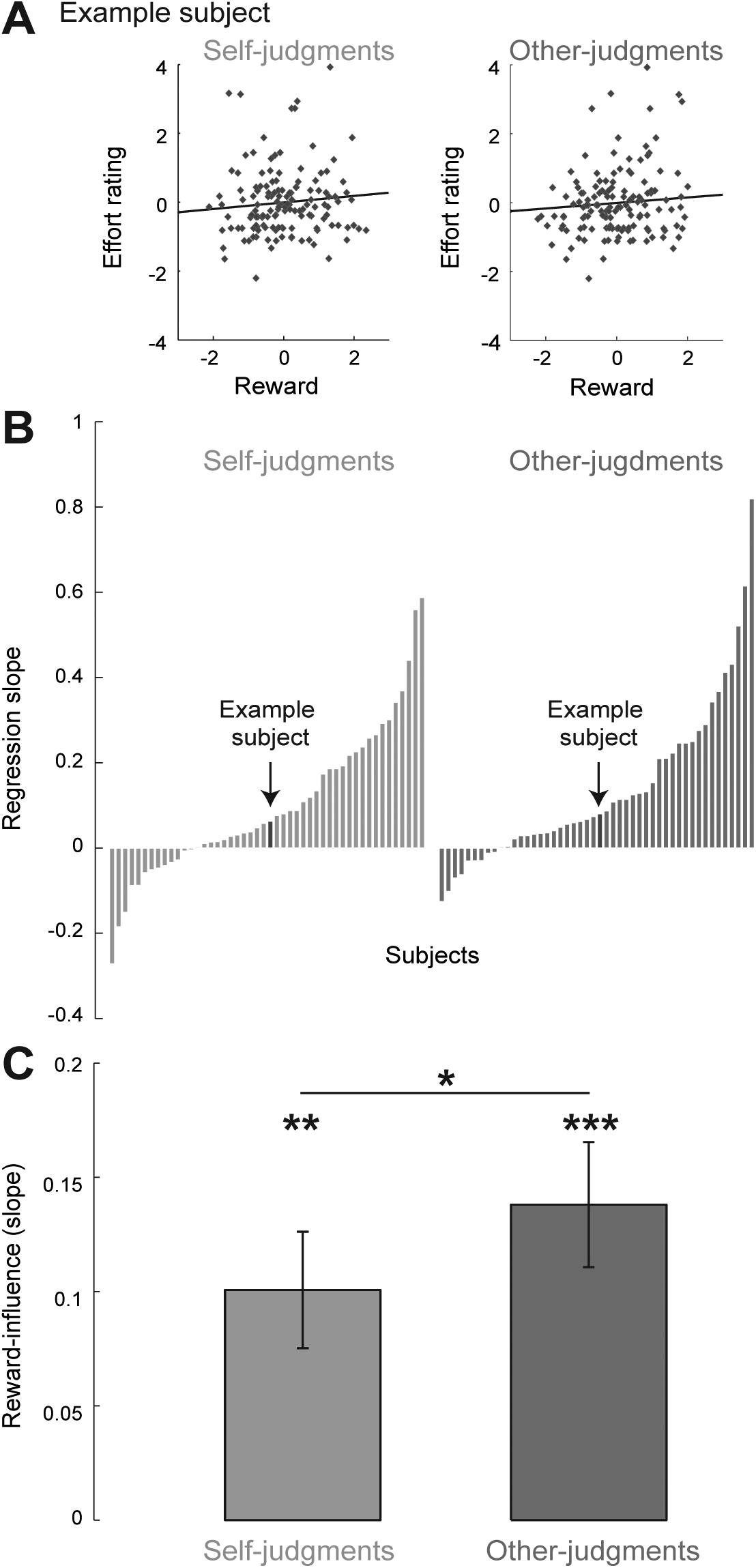
Influence of reward on effort judgments for self and others. For estimating the influence of reward on effort estimations, the trial-by-trial fluctuations in reward were used as predictors in a linear regression for estimated effort, separately for self- and other-judgments of each subject. **(A)** Data of one representative subject is shown as scatter plot and linear regression between trial by trial variation in reward and effort for self- (left) and other-judgments (right). Trial by trial reward magnitude and effort rating were z-scored within each difficulty level, so that values represent variation in comparison to the average of the difficulty level. **(B)** The individual (standardized) regression slopes for self and other ratings are presented. The example subject presented in (A) is indicated by the arrows. **(C)** Group average of slopes ± standard errors are presented (N=48). One-sample signed test and Wilcoxon signed rank test, two-sided ^∗^p<0.05, ^∗∗^p<0.01, ^∗∗∗^p<0.001

Having established an influence of reward magnitude on self- and other-judgments of effort, we next investigated how rewards were integrated with other information sources related to the exerted effort to form the final effort ratings. To this end, we compared predictions of five computational models with the behavioural data. The focus of these models was on how effort estimations, based on performance-related information (E_*p*_) and on reward information (E_*r*_), are combined to form the final effort rating (**Figure 3A**).

**Figure 3.**
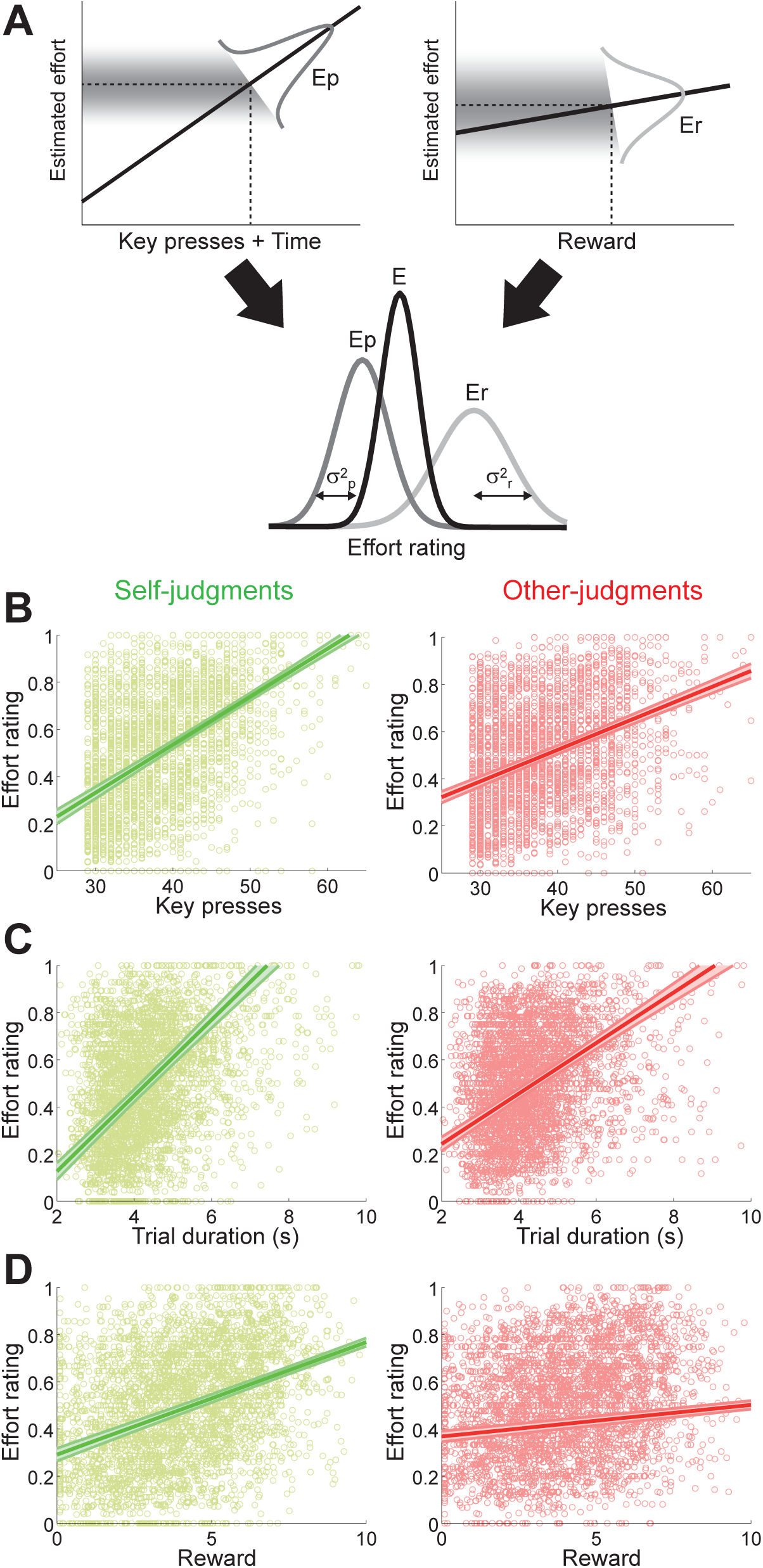
Different information sources are integrated to form a final effort rating. **(A)** Two types of information can be combined to derive the final effort estimation (E): E_p_ is the effort estimation based on performance-related information (key presses and trial duration), whereas E_r_ is the effort estimation based on reward-related information. On every trial, E_p_ and E_r_ depend on the specific amount of key presses, time and reward. Both estimates will differ by some degree. According to Bayes optimal models, the influence of each signal on E depends on its reliability, represented by the width of the distributions 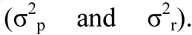
**(B)** Relation between the number of key presses and effort ratings for self- (left) and other-judgments (right), based on the trials in which effort was rated *before* the reward was revealed (30% of all trials). Scatter plots (light colours) show the corresponding data pulled over all 48 subjects. The regression line shows the average slope for all subjects ± the group standard error of the individual slopes. **(C)** Similar to (B) for the relation between trial duration and effort ratings. **(D)** Similar to (B) for the relation between reward magnitude and effort ratings. Note that this relation effectively captures the correlation between reward and effortrating built-in by the task design: Since trials that are more difficult were associated with higher rewards and these trials were subjectively rated with higher exerted efforts, reward magnitude carried information about the subjective estimations of exerted effort in these trials.

The exerted effort in this task can be estimated based on performance-related information such as the amount of key presses and the duration of a given trial. Those quantities were observable for subjects in every trial, both for self- and other-judgments. Assuming that participants’ subjective effort estimations show some correspondence with performance-related information, it seems plausible that subjective effort estimations should be based (at least to some degree) on the amount of executed key presses and the elapsed time of a trial (as shown in **Figure 3B** and **Figure 3C**). We can leverage this to deduce subjects’ presumed internal effort representations based on such observable quantities. Thus, we can define the expected effort estimation given performance-related information (key presses and elapsed time) as follows:

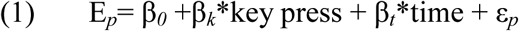

where β_*0*_, β_*k*_ and β_*t*_ and were fitted to the trials in which the effort was rated *before* the reward was revealed (30% of all trials), giving an unbiased estimation of effort judgments.

Moreover, based on the task design, rewards were contingent on exerted effort. Therefore, effort judgments can be predicted based on reward. We can define the expected effort estimation given reward (E_*r*_) as follows:

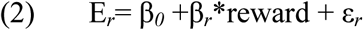

Analogous to Eq(1), we derived β_*0*_ and β_*r*_ from the trials in which the effort was rated *before* the reward was revealed. We then applied β_*k*_ and β_*t*_ as well as β_*r*_ to the trials where effort was rated *after* the reward was revealed, to get a trial-by-trial estimation of subjects’ internal effort representation based on either performance related information (E*_p_*) or reward information (E*_r_*) only.

Both effort estimations, E_*r*_ and E_*p*_, are associated with some uncertainty, representing how well exerted effort can be inferred from those different sources. The uncertainty associated with E_*r*_ and E_*p*_ can be formalized as the mean squared error:

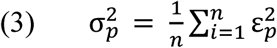

and

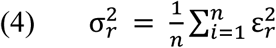

In each trial, E_*r*_ and E_*p*_ can differ by some amount (ΔE ≠ 0). We consider several models of how these two informational sources might be integrated into the actual effort rating E.

### Model 1

(performance only model) assumes the actual effort rating is only based on the E_*p*_ without taking reward information into account. Thus, the E*_p_* would directly result in the actual effort rating E:

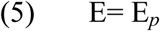

### Model 2

(reward only model), in contrast, only relies on reward information (E_*r*_) and completely ignores performance information:

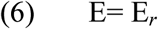

### Model 3

(simple average model) assumes that both sources of information are considered and that E is an equally weighted average of E_*r*_ and E_*p*_:

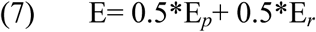

So far, the models did not take the reliability of the information into account. However, **Model 4** (Bayesian average model) assumes that the variance of the different sources of information is used in order to weight E_*p*_ and E_*r*_ in a Bayes optimal way (see **Figure 3A**):

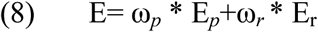

where
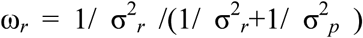
and ω_*p*_ =1-ω_*r*_ The variances
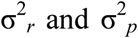 explained above (Eq(3) and Eq(4)) and the weights are directly derived from variances.

**Figure 4.**
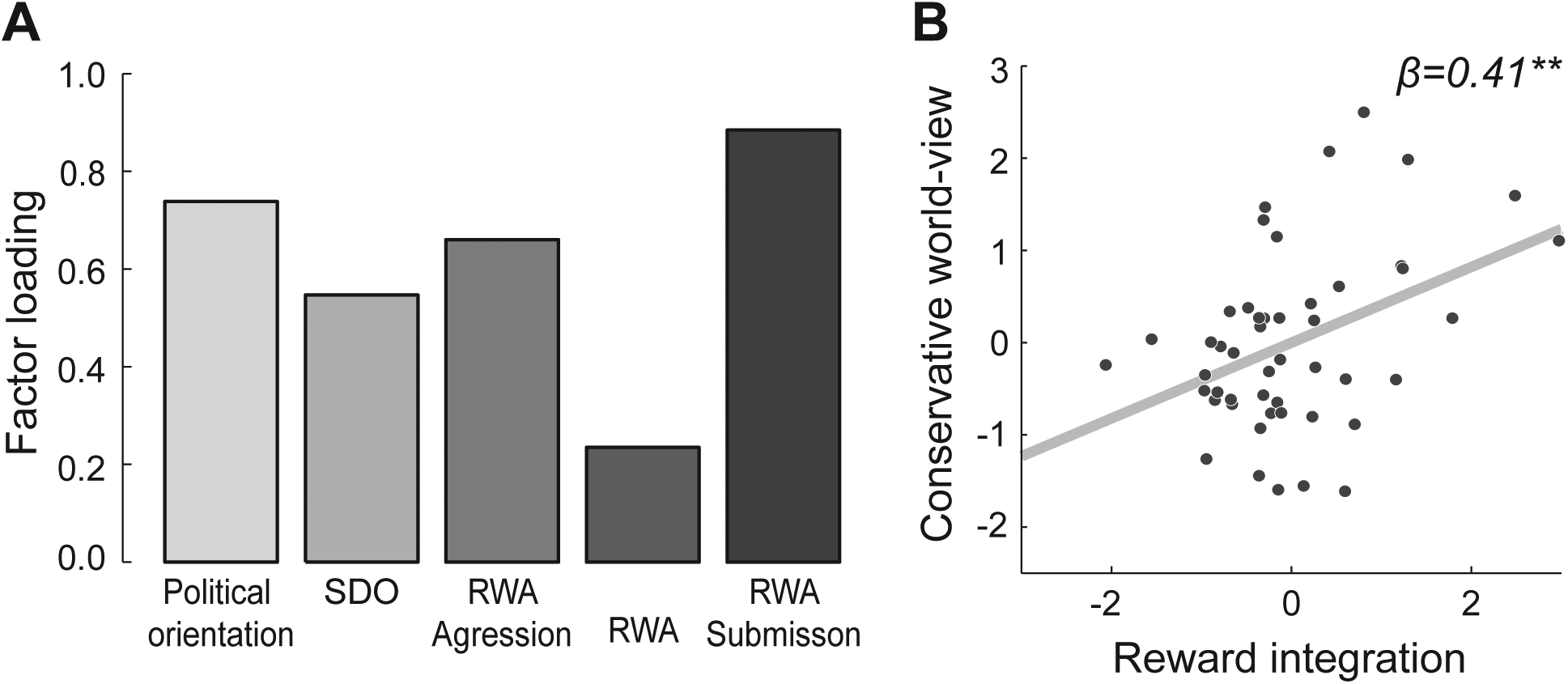
Conservative world-views (factor scores) are positively correlated with the amount of reward integration into effort estimation. (A) A factor analysis over multiple questionnaires revealed that one factor (based on scree-test) which can be interpreted as conservatism is explaining the interrelation between these scales. SDO: social dominance orientation; RWA: right-wing authoritarianism. (B) Stronger influence of rewards on effort estimations is positively associated with conservative world-view. The sum of the slopes for self- and other-judgments (slope_self_+slope_other_) is used as indicator for reward integration. Since we controlled for empathy and perspective taking in the regression analysis, we present values of reward integration from which we regressed out influences of these covariates (N=48). Linearregression, two-sided ^∗∗^p<0.01

Moreover, it might be possible that E_*r*_ and *E_p_* are combined, but with a different weighting than predicted by any of the previous models. Thus, in **Model 5** (flexible weighting model) we set the weighting parameter ω_*r*_ as a free parameter and ω_*p*_=1-ω_*r*_.

The model evaluation was done by computing the maximum-likelihood fits to the trial-to-trial data of individual subjects, separately for self- and other-judgments. The quality of fits was compared by the Bayesian Information Criterion (BIC; 25, 26).

The model comparison revealed that for both, self- and other-judgments, the Bayesian average model had the lowest BIC (**Table 1**), showing that this model provides the most accurate and parsimonious account of the data. For self-judgments the second best model was the flexible weighting model, whereas it was the simple average model for other-judgments.

**Table 1.**
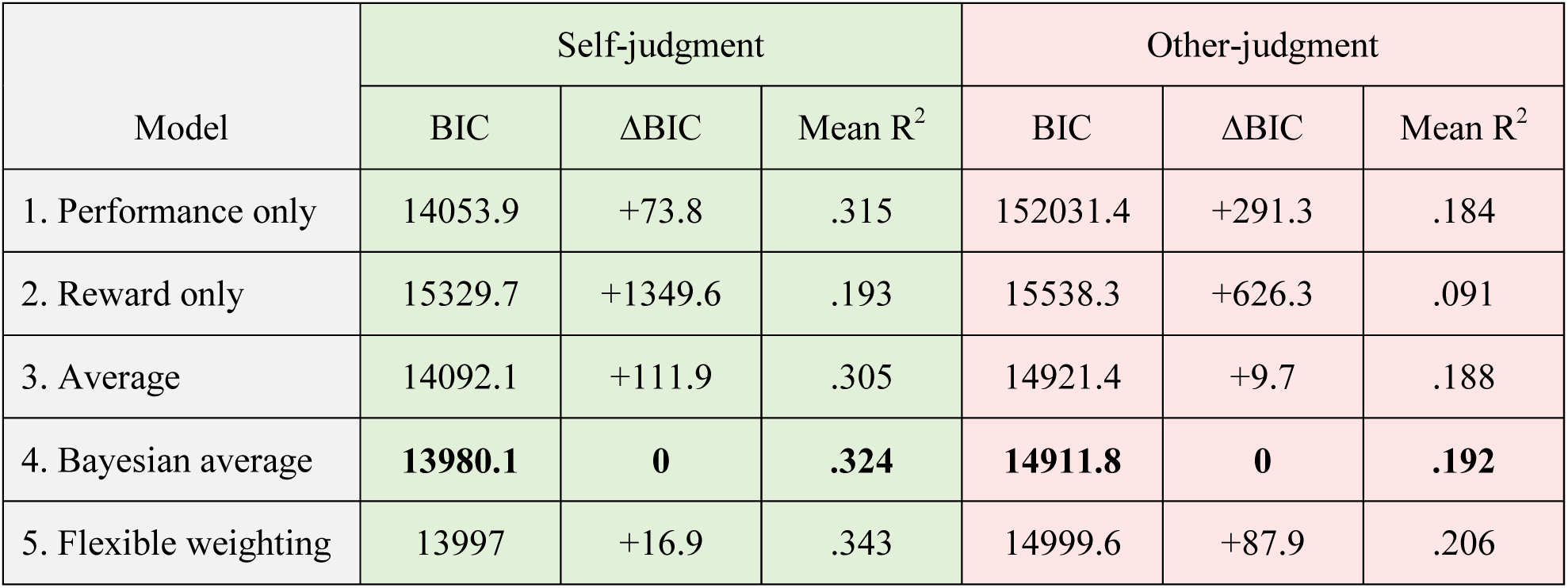
Results of model comparison. BIC: Bayesian Information Criterion. BIC values are summed across 48 subjects. The Bayesian average model (bold font) had the lowest BIC score and is therefore ranked highest by the model comparison. ΔBIC Difference in BIC of each model minus the BIC of the best fitting model (BIC_model_-BIC_best_).

These results provide additional evidence that reward information is integrated when forming effort estimations about self and others. Moreover, subjects integrate different sources of information weighted by their reliability, as predicted by Bayesian cue integration. Therefore, the integration of rewards into final effort ratings appears optimal in terms of information usage as rewards are combined with performance-related information based on the reliability with which each information source can be used to infer the exerted effort.

After having demonstrated that reward influences the estimation of exerted physical effort of self and others, we asked whether this behavioural effect showed any meaningful relationship with broader social attitudes. We hypothesized that reward-biased effort estimations, could be associated with acceptance of economic inequalities and reluctance to social change as these attitudes might represent a prior assumption about strong links between effort and reward/wealth. In order to probe a relationship between our behavioural effects and broader social attitudes, we assessed several questionnaires related to conservative world-views (27, 28; see Methods for detailed description ) and conducted a factor analysis on these scales using maximum-likelihood estimation. Based on the scree-test (29) we identified a sharp drop in eigenvalue after the first component (see **Suppl. Figure S2**), indicating that one factor, explaining 76% of the variance, was the best and most parsimonious solution for the covariance structure of the questionnaire data. We interpreted this factor as “*conservative world*-*views*” based on the factor loadings (**Figure 4A**). This factor represented acceptance/support of economic inequality, reluctance to social change and obedience to established authorities.

We extracted factor scores for every subject to relate those with our behavioural effect of reward integration into effort judgments. Due to the high correlation between reward integration for self- and other-judgments (cf. **Suppl. Figure S1**), we used the sum of both effects (from the model free analysis: slope_self_+slope_other_, see **Figure 2**) as indicator for the extent to which subjects were biased in their effort estimations by rewards.

In line with our hypothesis, subjects with stronger influence of reward magnitude on effort estimations displayed more conservative world-views (β=0.41, p=0.005, see **Figure 4B**), underlining an interesting link between this behavioural effect and social attitudes. Importantly, this association between conservatism and reward influence could also be shown when separately considering self-judgments (β=0.32, p=0.03) and other-judgments (β=0.44, p=0.002). It is remarkable that an association between a low-level behavioural laboratory task and high-level social attitudes could be found. In particular, the size of this effect is surprising since the influence of reward explained the variance of conservative attitudes to a similar extent as well-established socio-demographic predictors such as age, gender, education or income have been shown to do (30-32). Further analysis on whether the increased integration of reward for the conservative group is an irrational bias or is in line with Bayes optimal cue integration is shown in **Supplementary Information**.

## Discussion

Using retrospective evaluations of physical effort during sensorimotor task with monetary rewards, we found judgments about own and other people’s exerted effort to be influenced by reward magnitude, with more pronounced effects for other-judgments. Computational modelling revealed that both judgment types were constructed in a Bayes optimal manner, weighting performance related criteria and reward information by their reliability (inverse of variance). Furthermore, the extent to which individuals were influenced by rewards when judging efforts was positively correlated with conservative world-views.

Both self- and other-effort judgments were influenced by received rewards, showing that this effect is not an instance of self-serving bias but instead represents a general cognitive process of information usage. Importantly, the influence of rewards was even stronger when judging someone else’s effort. This finding is contrary to predictions from attribution theory that success (i.e. high reward) would be more strongly attributed to internal factors (i.e. exerted physical effort) for self-judgments compared to other-judgments (13). In contrast, other studies have found that people incorporate outcomes less when making judgments about themselves as compared to others (33), which is in line with our findings. In those studies the differential incorporation of outcomes for judging own and others’ behaviour has been described as an *introspection illusion* (33-35) suggesting that people put too much weight on subjective internal information when evaluating themselves, deviating from optimality. Our results however do not support the interpretation that people necessarily weigh internal and external information suboptimally during self-assessment. Using computational modelling we showed that people weigh internal (subjective representations of exertion level based on key presses and trial duration) and external (reward) information in a statistically optimal fashion where the weight of each information source is determined by its reliability. Furthermore, from Bayes optimal integration of information, it follows that subjects should incorporate rewards more when judging other people’s effort because they have less reliable estimates of others’ performance-related criteria.

The weighing of different relevant information sources by their reliability is the optimal way of combining information. In such a scheme, increased incorporation of reward information occurs when effort estimates based on performance criteria are less reliable. We propose that inferring the precision of internal representations of effort based on the actual physical exertion level (i.e. amount of key presses and elapsed time) is a form of metacognitive ability (36) enabling humans to accurately monitor and report their past mental states. It would be interesting to probe whether an improvement of this metacognitive ability (e.g. by training) might result in a reduced incorporation of reward information when rating one’s own effort, as would be predicted by the Bayesian framework. Furthermore, it is known that human brain is extremely sensitive to reward acquisition (37), and decision making agents are irrationally biased to repeat choices that recently led to real reward outcomes even when such outcomes were dissociated from long-term informational value (38). Therefore, it would be important to investigate if the receipt of a real monetary outcome such as used in our study has stronger influence than would be explained by the purely informational content of the reward feedback (or any other abstract task-contingent feedback).

We suggest that conservative people show stronger influences of rewards on effort estimations because they might have the prior belief that those two quantities are strongly related. However, integrating reward information into effort ratings can also be seen as a form of uncertainty reduction (39, 40). Interestingly, there is a solid body of literature suggesting uncertainty aversion as one of the core psychological foundations of conservative attitudes (see 22 for an extensive review). The association of conservative world-views with the increased reward integration might reflect additional information seeking or a pursuit of deterministic rationalization. It might be promising to revisit those existing psychological findings by using a computational approach, to see whether such findings could be accommodated within the Bayesian framework.

Independent of the underlying reason, more conservative people were more strongly influenced by rewards when judging their own and especially other people’s effort. We surmise that in some situations, such exaggerated influence of reward can lead to a biased perception of people’s exerted effort. Our results suggest that people with conservative views might judge other people’s effort more through the received gains, indicating that in everyday life they might adhere to the view that people with low income have also exerted less effort, and conversely, prosperous people deserve their wealth since they have worked harder, perhaps ignoring other contributing factors. A fine-grained understanding of the cognitive building blocks contributing to societal attitudes holds the potential for counteracting such biases and unfounded beliefs.

In summary, we have shown that retrospective decisions on the amount of exerted physical effort, both for self as well as for the others, are affected by the obtained rewards. While reward magnitudes are often explicit in naturalistic environments, effort demands are rarely explicit and are usually learned by trial and error in the course of successive actions (41). An important direction for future research is to determine how retrospective evaluations regarding the internal states and the state of the environment could in turn influence prospective value- and effort-based decisions, thus relating the effects we observed to their potential adaptive role in shaping human choices in individual and social contexts.

## Materials and Methods

### Participants

The final sample included 48 subjects (27 females, age 20-37 years, mean: 25.8 years). Ten subjects were excluded from the original sample (N=58) as there was indication of random ratings of effort (but we note that when not excluding any participants the results are qualitative similar, see **Supplementary Information** for results of the full sample). Task difficulty was the experimental factor controlling required effort, thus subjects should show an association between task difficulty and their effort ratings when performing the task properly, and those for whom effort ratings were not significantly influenced by task difficulty were excluded (i.e. no significant effect of a trial-by-trial linear regression between task difficulty and effort ratings, see **Suppl. Figure S3**). Participants were paid 8 Euro per hour, plus a bonus for their performance in the experiment, resulting in a total mean payment of 20 ± 5 Euro. All subjects gave written consent about their attendance. The study was approved by the local ethics committee of University Medicine Göttingen, proposal number **15/7/15**.

### Experimental design

The behavioural experiment was adapted from Pooresmaeili, Wannig and Dolan (17) in order to test two subjects simultaneously. Participants performed the task in pairs, interleaved with a partner. In half of the trials participants performed the task themselves and rated their own effort (self-judgment). In the other half, participants rated the other person’s effort (other-judgment) after watching them performing the task. Each participant in a pair performed 200 trials resulting in a total of 400 trials which were grouped into 10 experimental blocks (each containing 40 trials).

For more details on the task and stimuli, please see **Supplementary Information**.

### Questionnaires

Participants completed questionnaires about social attitudes and political orientation. We used a political orientation self-report (on a scale from 0=”very liberal” to 100=”very conservative”) to assess explicit conservative orientation. As indirect indicators for conservatism we also assessed social dominance orientation (27), reflecting a general preference for hierarchical intergroup relations, and right-wing authoritarianism (28), measuring authoritarian submission, authoritarian aggression, and conventionalism (42). Since we were interested in the underlying factor linking these scales, we conducted a factor analysis over these scales, using a maximum-likelihood estimation. The number of factors was based on the Scree-test (Cattell, 1966; see **Suppl. Figure S2**), indicating that one factor was sufficient to explain the interrelation between these questionnaires. Factor scores of the first factor were extracted to relate those to behavioural measures.

As conservatism, especially right-wing authoritarianism and social dominance orientation, were reported to correlate negatively with empathy (27, 43), and empathy could arguably have an impact on our behavioural effects of interest, we assessed empathy and perspective-taking as covariates (44).

### Statistical analysis

All presented results were tested two-tailed. Results from regression analysis are always reported as standardized betas. To investigate the relationship between conservative world-views and influence of rewards on effort estimations linear regression analysis was used, controlling for empathy and perspective-taking as covariates. The sum of slopes for self- and other-judgments between reward magnitude and effort estimations was used as predictor and the factor scores from the factor analysis were used as indicator for conservative world-views.

For more details of the statistical analysis see **Supplementary Information**.

## Acknowledgements

We thank Aurel Wannig and Thomas Schultze-Gerlach for their valuable comments on the manuscript. This work was supported by a seed fund grant from Leibniz ScienceCampus *Primate Cognition*, Göttingen, Germany to IK and AP. AP was supported by an ERC Starting Grant (716846, *Rewarded Perception*). The Wellcome Centre for Human Neuroimaging is supported by core funding from the Wellcome Trust (203147/Z/16/Z).

## Authors’ contributions

MR, IK and AP conceptualized and designed the task. MR and UT chose and devised the online questionnaires. MR and FP conducted the experiments. MR and CS analysed the data. MR, IK and AP wrote the manuscript.

## Data availability

The datasets generated and analysed in the current study are available from the corresponding author upon request, and will be uploaded to Open Science Framework data repository (https://osf.io/).

## Supplementary Information

### Experimental design

The behavioural experiment was adapted from Pooresmaeili, Wannig and Dolan (17) in order to test two subjects simultaneously, and lasted around 2 hours. Participants pressed two keys of the keyboard, as fast as possible, to push a ball up a virtual ramp before rating the exerted effort. Trials were presented interleaved between both subjects. In half of the trials participants performed the task themselves and rated their own effort (self-judgment). In the other half, participants rated the other person’s effort (other-judgment) after watching them performing the task.

While the participants tried to push the ball up the ramp with fast alternating key presses, the ball rolled back by a constant amount during each frame of the display, simulating gravity force. This gravity force differed pseudo-randomly between trials and created the six difficulty levels of the task (individually adjusted for each participant, see below). If the participants managed to push the ball up the ramp within 10 seconds they were rewarded for this successful trial (only the participant who performed the trial was rewarded). The amount of reward was contingent on the task difficulty and the actual reward value was drawn from Gaussian distributions with means ranging from 1.5 to 6.5 cents and standard deviation of 1.2 cents. In most of the trials (70%) the reward was presented before subjects rated their effort and in some of the trials (30%) the reward was presented after participants had judged the effort. Trials in which reward was shown after the estimation of effort served as a reference, since in those trials no influence of reward was present.

The subjects were seated inside the same room, next to each other, separated by a partition wall. Each subject in a pair performed 200 trials resulting in a total of 400 trials which were grouped into 10 experimental blocks (each containing 40 trials). The subjects saw on their screen how the ball was moved up the ramp, independently of whether they performed the task themselves or whether the other person was playing. Moreover, subjects wore headphones and each key press was accompanied by a beep tone. Hence, subjects had visual and auditory information about the key presses. In trials where the participants conducted the task themselves they judged their own exerted effort and in trials when the other person performed the task, they estimated the effort that the other person had performed. Subjects did not see how the other person rated the effort, in order to prevent reciprocal influences.

### Stimuli and Task

The stimulus presentation and registration of behavioural responses was controlled by one personal computer. Stimuli were produced with MATLAB and the Psychophysics Toolbox. Each trial consisted of a stimulus (ball and ramp), a reward, and an effort rating display all shown on a black background (a trial timeline is shown in **Figure 1A**). The stimulus display contained the ball (radius: three visual degrees), initially at the starting part of the ramp (ramp length: 19 visual degrees; both ball and ramp had a light grey colour). The ball was displaced up the ramp with consecutive alternate key presses (left and right arrow keys) until it reached the upper plateau. Each key press resulted in a constant amount of displacement (0.87 visual degrees per key press) and was counteracted by a gravity force of variable strength that displaced the ball backward. Moreover, each key press caused a beep tone which was played on the headphones of both participants. To determine the levels of used gravity force individually for each subject, at the beginning of the experiment we asked each subject to push the ball up the ramp by pressing both keys alternately and consecutively as fast as they possibly could. To motivate the participants, they were told that they would receive 1 Euro bonus if they managed to push the ball up the ramp. Ninety percent of the gravity force necessary to counteract the maximum number of key presses in a limited time (10 s) determined the maximum gravity force used in the experiment. This estimation was done for each participant alone; the other participant in a pair waited outside the room. Based on this individualized estimate, six equally spaced gravity levels were defined and used in the experiment. A trial was aborted if key presses did not occur fast enough (maximum pause allowed was 2 s). If participants were able to successfully push the ball all of the way up, they received a monetary reward, with the amount contingent on task difficulty (gravity force level). Reward magnitude was defined based on six Gaussians with means of 1.5, 2.5, 3.5, 4.5, 5.5, and 6.5 cents with a SD of 1.2 cents. Reward display consisted of a pie chart that depicted subjects’ reward as a proportion of maximum reward possible and a number that showed the reward in Arabic numerals. Effort rating display consisted of a slider, and participants were instructed to set the slider at a position that represented their experienced effort during a trial proportionate to the maximum effort they could experience during the experiment. In 70% of trials, the reward display was shown immediately after the stimulus display, whereas in 30% of trials it occurred after the rating of effort. Before the actual experiment started, participants conducted 36 training trials (which were not analysed) to get familiar with the experiment and to learn about the effort-reward contingency. Because task difficulty and reward were the only parameters that varied across trials, we intended to distract participants from the main purpose of this experiment to prevent ad hoc strategies of relating reward and effort. For this reason, we introduced a second task into the paradigm where subjects were asked to report whether they had seen a brief (21 ms or three frames) colour change on the ball (to green, red, or blue), which occurred in 50% of the trials. This question was asked randomly in 20% of trials.

### Statistical analysis

Outlier values of effort ratings (3 standard deviations above or below the participant’s mean effort) were removed from the analysis, since some subjects reported in the debriefing to have occasionally rated extreme effort values to test their assumptions about the aim of the experiment (e.g. whether they could influence the received reward by their effort ratings). However, when not excluding any effort ratings from the analysis the results are qualitatively similar (see **below** for results without any excluded effort ratings).

For the analysis of the relationship between trial-to-trial variation in reward magnitude and effort ratings, both were z-scored within difficulty levels, resulting in values representing variation in reward magnitude and effort rating with respect to the average of the corresponding difficulty levels. After having created these within difficulty variations we pooled them across all difficulty levels and conducted a robust linear regression (robust regression toolbox in MATLAB) with reward as predictor and effort rating as dependent variable.

For comparison of the computational models, each model was fitted to participants’ z-scored effort ratings, separately for self- and other-judgments, and BIC scores were calculated. For model comparison the sum of individual BIC scores was used.

To investigate the relationship between conservative world-views and influence of rewards on effort estimations linear regression analysis was used. Factor scores of the first factor from the factor analysis were used as indicator for conservative world-views. The sum of slopes for self- and other-judgments between reward magnitude and effort estimations was used as indicator for the influence of rewards on effort estimations. The influence of rewards on effort estimations was used as predictor and conservative-world views as dependent variable in the linear regression analysis, controlling for empathy and perspective-taking as covariates.

To investigate whether conservative world-views were related to weights of reward-information (ω_*r*_) predicted by the Bayesian average model, we conducted linear regression analysis with conservative world-views as predictor and ω_*r*_ as dependent variable. To further investigate whether this association was driven by less reliable effort estimations based on performance information we conducted a linear regression between conservative world-views and σ^2^_*p*_.

To investigate whether there is an association between conservative attitudes and the extent to which subjects’ behaviour was in line with Bayesian cue integration we calculated the BIC-weight of the Bayesian average model for every subject as follows:

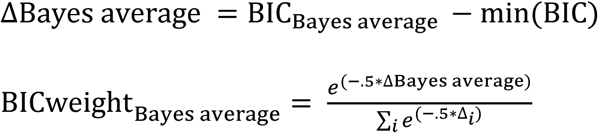

The Bayesian weights of all of the models in a set sum up to 1 and indicate the probability for each model to be the best model for the data.

We then calculated a linear regression with conservative world-views as independent variable and BICweight_Bayes average_ as dependent variable.

### Bayes optimality in people with low and high conservatism scores

Since we have shown that the integration of rewards into effort estimations can be optimal with respect to the use of available information, we asked whether the increased integration of reward information for conservatives is an irrational bias or whether it is in line with Bayes optimal cue integration. One possibility is that conservative group had access to less reliable effort estimations based on the available performance information (key presses and time), in which case it would be beneficial to weight the reward information stronger. Alternatively, conservatism could lead to integration of reward information above the extent that would be Bayes optimal.

To this end, we investigated whether conservatism was associated with the weights of reward information (ω_r_) predicted by the Bayesian averaging model. There was a significant association between ω_r_ and conservative world-views for self- (β=0.3, p=0.04) but not for other-judgments (β=0.21, p=0.15), indicating that the increased incorporating of rewards into effort estimations for self-judgments is in line with Bayesian cue integration. The increased weight for reward information was marginally driven by less reliable (self-) effort estimations based on performance, indicated by a trend towards positive association between uncertainty σ^2^_*p*_ and conservative world-views (β=0.27, p=0.076). Since the increased reward integration within the conservative group for other-judgments was not correlated with the weights of reward information (ω_r_ predicted by the Bayesian model, we investigated whether conservative people deviated from Bayes optimality for those judgments. To this end, we conducted the model comparison for other-judgments separately for people with low and high conservatism scores (based on a median-split). The group with low conservatism scores was best described by the Bayesian average model (difference in BIC relative to next best model: 12.0), however the group with high conservatism scores was best described by the simple average model (BIC=7253.6) followed by the Bayesian average model (BIC=7256). In line with this, the extent to which each subject’s other-judgments were captured by the Bayesian average model (BICweight, see Methods for a detailed description) was negatively correlated with conservative attitudes, a trend that did not reach statistical significance (β=-0.26, p=0.078). These results suggest that the increased incorporation of rewards by the conservative group is in line with Bayes optimality for self-judgments, whereas when judging someone else’s effort people with high conservative scores deviate from Bayes optimality. We note however that results regarding Bayes optimality in conservatives are relatively weak and thus might depend to some extent on the exact formulation of the used models and the investigated model space.

### Results without applying exclusion criteria

In the main text we report results in which 10 subjects were excluded due to random effort ratings (their effort ratings were not significantly influenced by task difficulty, based on a linear regression with effort ratings as dependent variable and task difficulty as predictor), as well as excluded effort ratings that were ± 3 standard deviations from the subject’s mean effort ratings. These exclusion criteria were applied to make sure that only effort ratings that were not randomly chosen contributed to the analysis. However, here we show that the results are qualitatively similar when not excluding any subjects (using the full sample of N=58) nor outlier effort ratings from the analysis.

Across subjects, there was a significant influence of reward variation on effort estimation for self-judgments (one-sample signed test, p =0.002) as well as for other-judgments (one-sample signed test, p =0.0001) and the effect of rewards on effort estimations was significantly stronger for other-than self-judgments (mean slopes of 0.12 and 0.09 respectively, Wilcoxon signed rank test, p =0.028). The amount of reward integration for self- and other-judgments was highly correlated across subjects (r=0.78, p<10^-12^).

The comparison of the computation models showed that the Bayesian average model had the lowest BIC for self- (BIC= 16974.8) and other-judgments (BIC= 17919.3), followed by the flexible weighting model for self-judgments (BIC= 17009.6) and the simple average model for other-judgments (BIC= 17934.5).

Regarding associations of the integration of reward into effort judgments and conservative world-views, there was a positive association with the sum of the self- and other-judgment effect (β=0.32, p=0.018). This association between conservatism and reward influence could also be shown when considering other-judgments (β=0.34, p=0.011) and a trend was present for the association for self-judgments (β=0.25, p=0.063).

With respect to whether or not the increased reward integration for conservatives was in line with Bayesian cue integration we found a significant association between the weights of reward information (ω_r_) predicted by the Bayesian averaging model and conservative world-views for self-judgments (β=0.33, p=0.012), but not for other-judgments (β=0.16, p=0.25). This increased weight for reward information seemed to be driven by less reliable (self-) effort estimations based on performance information, indicated by a trend level positive association between uncertainty σ^2^_*p*_ and conservative world-views (β=0.25, p=0.074). However, the picture about other-judgments was less clear. When conducting the model comparison for other-judgments separately for people with low and high conservatism scores (based on a median-split) the group with low conservatism scores was clearly best described by the Bayesian average model (difference in BIC relative to next best model: 13.2), however the group with high conservatism scores was also best described by Bayesian average model (BIC=7256) although it was only slightly better than the simple average model (difference in BIC: 1.96). There was a trend that the extent to which each subject’s other-judgments were captured by the Bayesian average model (BICweight) was negatively correlated with conservative attitudes (β=-0.21, p=0.11).

### Influence of reward on effort ratings, controlling for within-difficulty fluctuations in gravity (due to trial-specific timing of presentation software)

Our main analysis for investigating the influence of reward on effort ratings focussed on a regression between variations of reward magnitude and effort ratings within difficulty levels (**Figure 2**). The rationale of this analysis was based on the fact that for each difficulty rewards were drawn randomly and gravity (the velocity of the ball rolling backwards) was fixed. Therefore, reward magnitude should have been the only variable that could change across trials of the same difficulty level. However, due to differences in exact timing of the presentation software, even within each difficulty level there were slight fluctuations of the actual experienced gravity force. Although such within-difficulty fluctuations should theoretically only have added noise to the data (as they were unrelated to received reward magnitude) and therefore decreased the observed effect of rewards on effort ratings, we wished to ensure that our results were not influenced by these variations. Therefore, we conducted the same regression between reward magnitude and effort ratings as shown in **Figure 2**, but controlling for the within-difficulty variation of trial-by-trial gravity force as covariate. As expected, when controlling for the within-difficulty fluctuations of gravity all results stay qualitatively the same, if anything with more pronounced effects. There was a positive influence of reward on effort ratings both for self- (one-sample signed test, p <0.0001) and other-judgments (one-sample signed test, p <0.00001) and this effect was stronger for other-judgements than self-judgements (mean slopes of 0.14 and 0.1 respectively, Wilcoxon signed rank test, p =0.041). These results indicate that our findings were not driven by within-difficulty variations of gravity force.

**Supplementary Figure S1.**
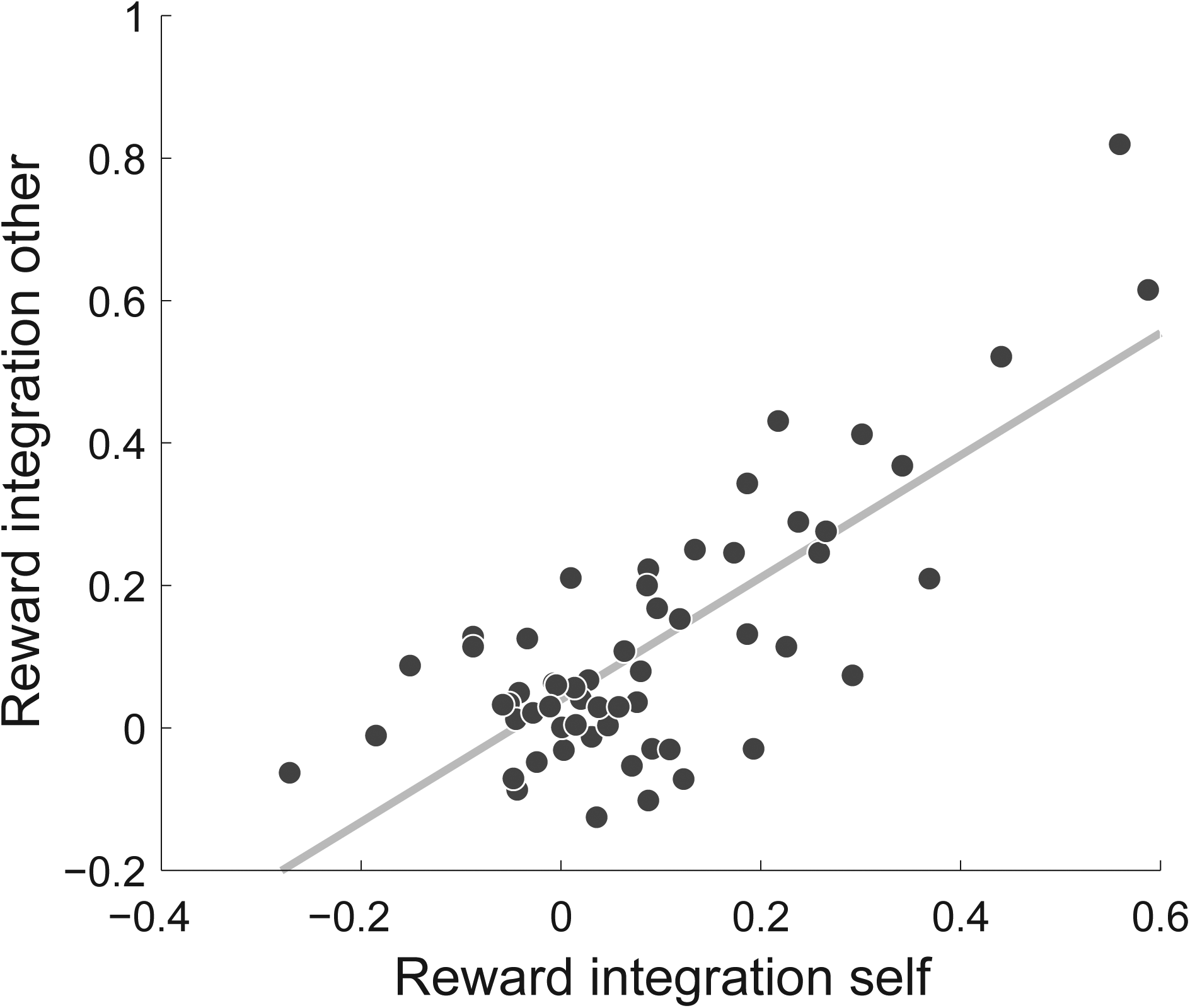
The amount of reward integration for self- and other-judgments was highly correlated across subjects. The standardized regression slopes between reward magnitude and effort ratings are used as indicator for reward integration. This measure was calculated separately for self- and other-judgments for every subject (N=48).

**Supplementary Figure S2.**
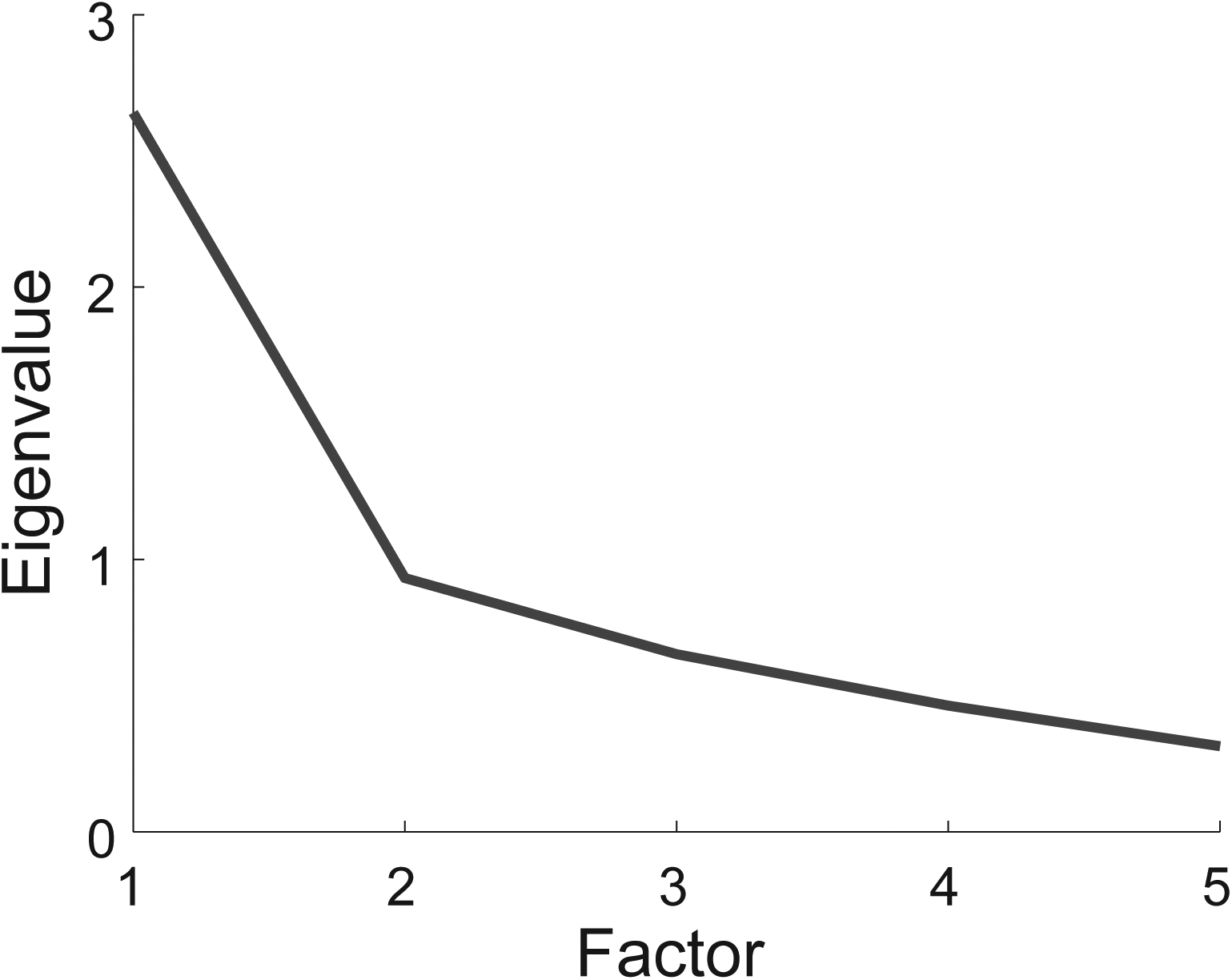
The scree plot shows that one factor is enough to capture the interrelation between several questionnaires used to assess conservative world-views. Eigenvalues for up to five possible components are presented. There is a sharp decline in eigenvalue from the first to the second factor, forming a clear “elbow”.

**Supplementary Figure S3.**
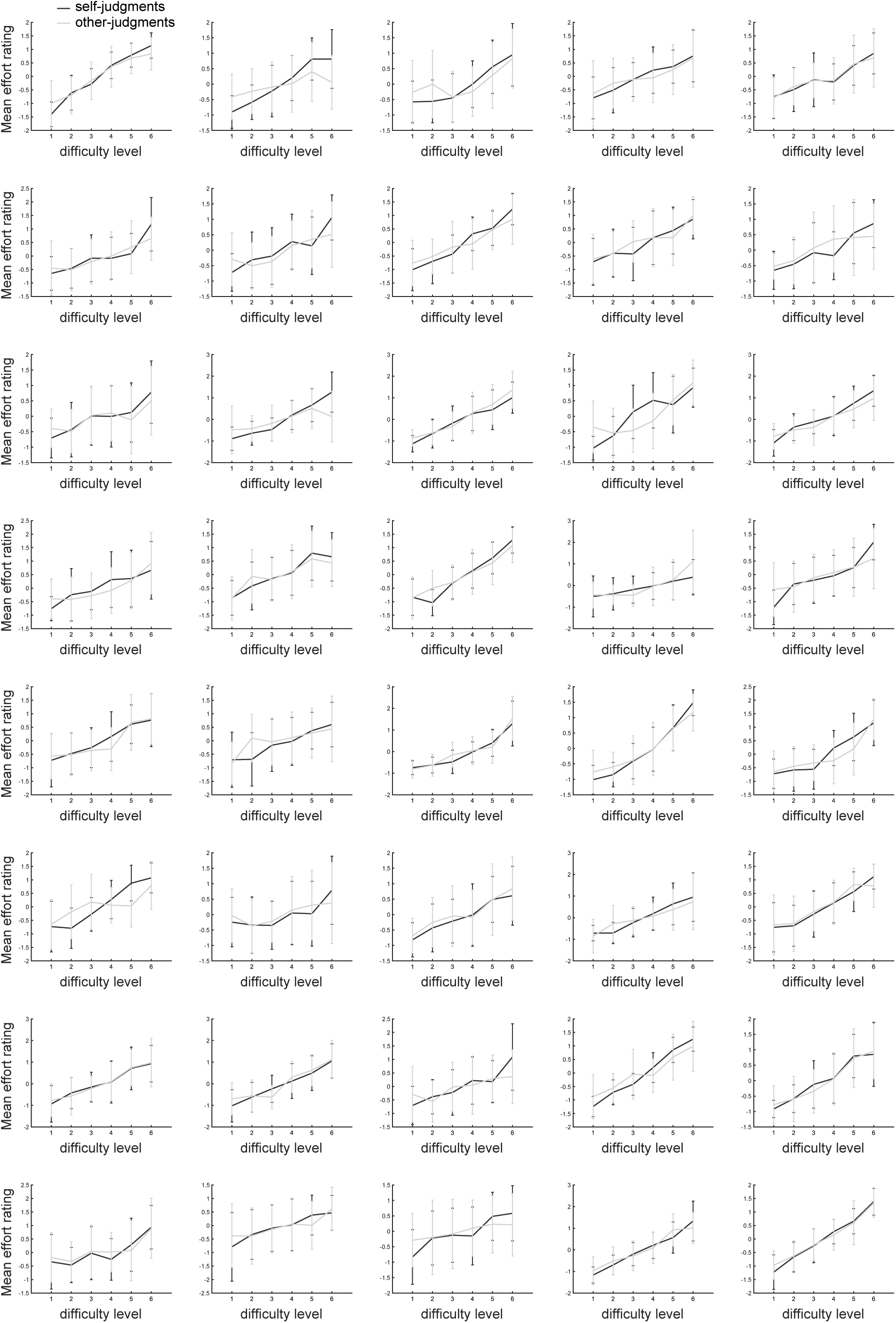
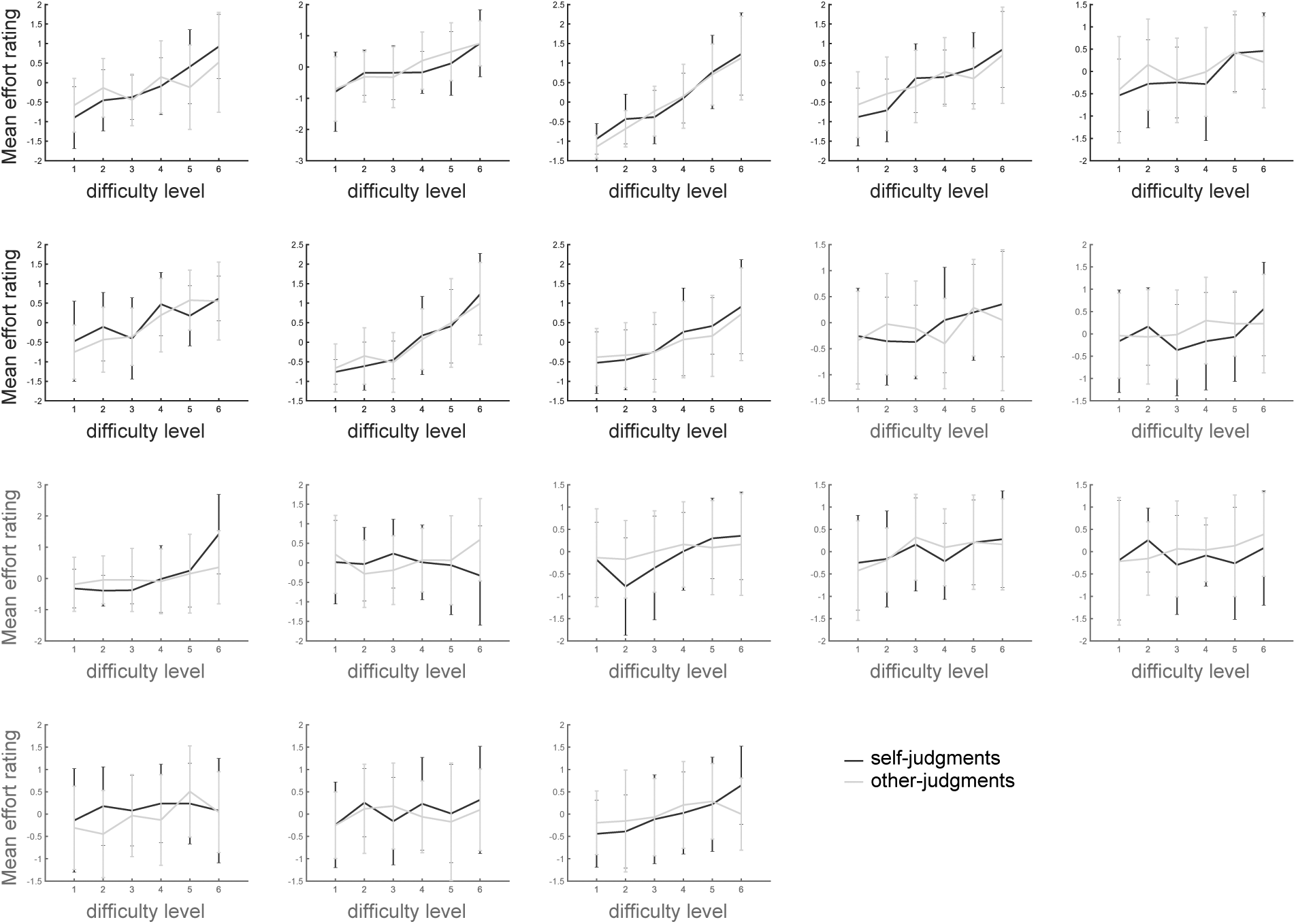
The psychometric plots showing the relationship between task difficulty and effort ratings for each subject, in trials where reward feedback preceded the effort rating. Mean effort ratings (± standard deviation) for each difficulty level are shown, separately for self- (blue curves) and other-judgments (green curves). Ten subjects that did not show a significant relationship between task difficulty and effort ratings were excluded from the final sample. Those excluded subjects are ordered at the bottom of the figure and marked by red coloured axes.

## References

1. Tjosvold D, Johnson DW, & Johnson RT (1981) Effect of partner’s effort and ability on liking for partner after failure on a cooperative task. The Journal of Psychology: Interdisciplinary and Applied 109(1): 147-152.

2. Fehr E & Gachter S (2002) Altruistic punishment in humans. Nature 415(6868): 137-140.

3. Fehr E & Gachter S (2000) Cooperation and Punishment in Public Goods Experiments. The American Economic Review 90(4):980-994.

4. Feldman S (1983) Economic Individualism and American Public Opinion. American Politics Quarterly 11(1):3-29.

5. Walton ME, Bannerman DM, Alterescu K, & Rushworth MF (2003) Functional specialization within medial frontal cortex of the anterior cingulate for evaluating effort-related decisions. J Neurosci 23(16):6475-6479.

6. Pessiglione M, et al. (2007) How the brain translates money into force: a neuroimaging study of subliminal motivation. Science 316(5826):904-906.

7. Rangel A, Camerer C, & Montague PR (2008) A framework for studying the neurobiology of value-based decision making. Nat Rev Neurosci 9(7):545-556.

8. Skvortsova V, Palminteri S, & Pessiglione M (2014) Learning to minimize efforts versus maximizing rewards: computational principles and neural correlates. J Neurosci 34(47):15621-15630.

9. Apps MA, Grima LL, Manohar S, & Husain M (2015) The role of cognitive effort in subjective reward devaluation and risky decision-making. Sci Rep 5:16880.

10. Klein-Flugge MC, Kennerley SW, Friston K, & Bestmann S (2016) Neural Signatures of Value Comparison in Human Cingulate Cortex during Decisions Requiring an Effort-Reward Trade-off. J Neurosci 36(39):10002-10015.

11. Morel P, Ulbrich P, & Gail A (2017) What makes a reach movement effortful? Physical effort discounting supports common minimization principles in decision making and motor control. PLoS Biol 15(6):e2001323.

12. Yeung N & Summerfield C (2012) Metacognition in human decision-making: confidence and error monitoring. Philosophical Transactions of the Royal Society B: Biological Sciences 367(1594): 1310-1321.

13. Weiner B (1972) Attribution Theory, Achievement Motivation, and the Educational Process. Review of Educational Research 42(2):203-215.

14. Aronson E & Mills J (1959) The effect of severity of initiation on liking for a group. The Journal of Abnormal and Social Psychology 59(2): 177-181.

15. Glimcher P (2014) Understanding the Hows and Whys of Decision-Making: From Expected Utility to Divisive Normalization. Cold Spring Harb Symp Quant Biol 79:169-176.

16. Tsetsos K, et al. (2016) Economic irrationality is optimal during noisy decision making. Proceedings of the National Academy of Sciences 113(11):3102.

17. Pooresmaeili A, Wannig A, & Dolan RJ (2015) Receipt of reward leads to altered estimation of effort. Proc Natl Acad Sci U S A 112(43): 13407-13410.

18. Mezulis AH, Abramson LY, Hyde JS, & Hankin BL (2004) Is there a universal positivity bias in attributions? A meta-analytic review of individual, developmental, and cultural differences in the self-serving attributional bias. Psychol Bull 130(5):711-747.

19. Ernst MO & Banks MS (2002) Humans integrate visual and haptic information in a statistically optimal fashion. Nature 415(6870):429-433.

20. Kording KP & Wolpert DM (2004) Bayesian integration in sensorimotor learning. Nature 427(6971):244-247.

21. Jost JT, Federico CM, & Napier JL (2009) Political ideology: its structure, functions, and elective affinities. Annu Rev Psychol 60:307-337.

22. Jost JT, Glaser J, Kruglanski AW, & Sulloway FJ (2003) Political conservatism as motivated social cognition. Psychol Bull 129(3):339-375.

23. Cozzarelli C, Wilkinson AV, & Tagler MJ (2001) Attitudes toward the poor and attributions for poverty. Journal of Social Issues 57(2):207-227.

24. Mccoy SK & Major B (2007) Priming meritocracy and the psychological justification of inequality. Journal of Experimental Social Psychology 43(3):341-351.

25. Burnham K & Anderson D (2002) Model selection and multimodel inference: a practical information-theoretic approach (Springer, New York).

26. Lewandowsky S & Farrell S (2010) Computational Modeling in Cognition: Principles and Practice (SAGE).

27. Pratto F, Sidanius J, Stallworth LM, & Malle BF (1994) Social dominance orientation: A personality variable predicting social and political attitudes. Journal of Personality and Social Psychology 67(4):741-763.

28. Funke F (2005) The Dimensionality of Right-Wing Authoritarianism: Lessons from the Dilemma between Theory and Measurement. Political Psychology 26(2): 195-218.

29. Cattell RB (1966) The Scree Test For The Number Of Factors. Multivariate Behav Res 1(2):245-276.

30. Ortoleva P & Snowberg E (2015) Overconfidence in Political Behavior. American Economic Review 105(2):504-535.

31. Sidanius J, Pratto F, & Rabinowitz JL (1994) Gender, Ethnic Status, and Ideological Asymmetry:A Social Dominance Interpretation. Journal of Cross-Cultural Psychology 25(2): 194-216.

32. Saucier G (2000) Isms and the structure of social attitudes. J Pers Soc Psychol 78(2):366-385.

33. Pronin E, Berger J, & Molouki S (2007) Alone in a crowd of sheep: asymmetric perceptions of conformity and their roots in an introspection illusion. J Pers Soc Psychol 92(4):585-595.

34. Pronin E (2007) Perception and misperception of bias in human judgment. Trends Cogn Sci 11(1):37-43.

35. Pronin E, Gilovich T, & Ross L (2004) Objectivity in the eye of the beholder: divergent perceptions of bias in self versus others. Psychol Rev 111(3):781-799.

36. Fleming SM, Dolan RJ, & Frith CD (2012) Metacognition: computation, biology and function. Philos Trans R Soc Lond B Biol Sci 367(1594): 1280-1286.

37. Vickery TJ, Chun MM, & Lee D (2011) Ubiquity and specificity of reinforcement signals throughout the human brain. Neuron 72(1): 166-177.

38. Scholl J, et al. (2015) The Good, the Bad, and the Irrelevant: Neural Mechanisms of Learning Real and Hypothetical Rewards and Effort. J Neurosci 35(32): 11233-11251.

39. Hasson U (2017) The neurobiology of uncertainty: implications for statistical learning. Philosophical Transactions of the Royal Society B: Biological Sciences 372(1711).

40. Hirsh JB, Mar RA, & Peterson JB (2012) Psychological entropy: a framework for understanding uncertainty-related anxiety. Psychol Rev 119(2):304-320.

41. Hauser TU, Eldar E, & Dolan RJ (2017) Separate mesocortical and mesolimbic pathways encode effort and reward learning signals. Proc Natl Acad Sci U S A 114(35):E7395-E7404.

42. Altemeyer B (1981) Right-wing authoritarianism. (University of Manitoba Press, Winnipeg, Canada).

43. Lippa R & Arad S (1999) Gender, personality, and prejudice: The display of authoritarianism and social dominance in interviews with college men and women. Journal of Research in Personality 33(4):463-493.

44. Maes J, Schmitt M, & Schmal A (1995) Gerechtigkeit als innerdeutsches Problem: Werthaltungen, Kontrollüberzeugungen, Freiheitsüberzeugungen, Drakonität, soziale Einstellungen, Empathie und protestantische Arbeitsethik als Kovariate (Univ., Fachbereich I, Psychologie, Trier).

